## Supplemental Figures for "Parahippocampal neurons encode task-relevant information for goal-directed navigation"

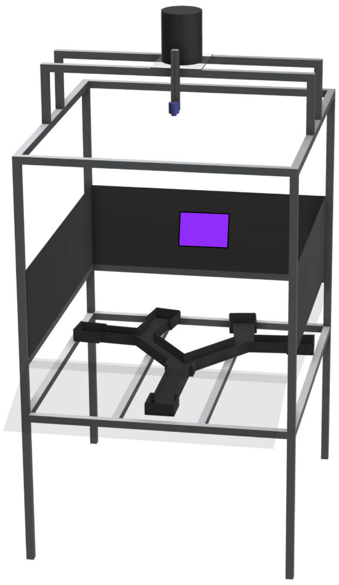

Supplemental Figure 1: Data collection and behavior apparatus. An aluminum frame housed both the open-field arena and Tree-Maze maze. The open-field "floors" were easily swapped in for open-field recordings, or taken out for Tree-Maze recordings. The height difference between the two was 0.35m, with all other peripheral cues staying constant across recording types. LED panel shown in purple at the back of the maze represents the "Right Cue".

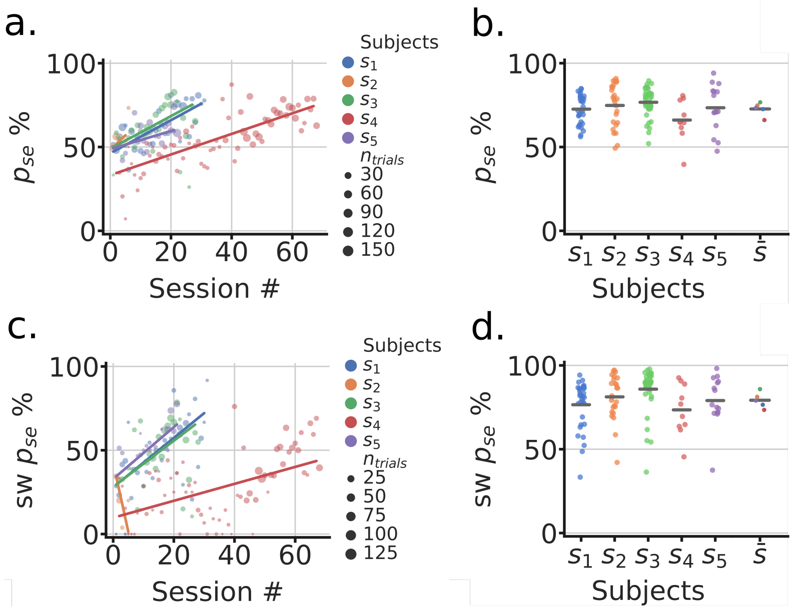


Supplemental Figure 2: Subject learning and post-surgery behavior. **a.** Tree-Maze performance by training session until performance criteria was reached (# trials=80, $p_{se} = 75\%$). Each dot corresponds to the performance on a given session, with colors identifying subjects and dot size indicating the number of trials in the session. Lines correspond to least squares regression by subject. **b.** Subject performance on the Tree-Maze task, post-surgery (Same as Fig.1d). Neural recordings were performed during all of these sessions. **c,d**. Same as (**a**) and (**d**), respectively, but for performance on switch trials.

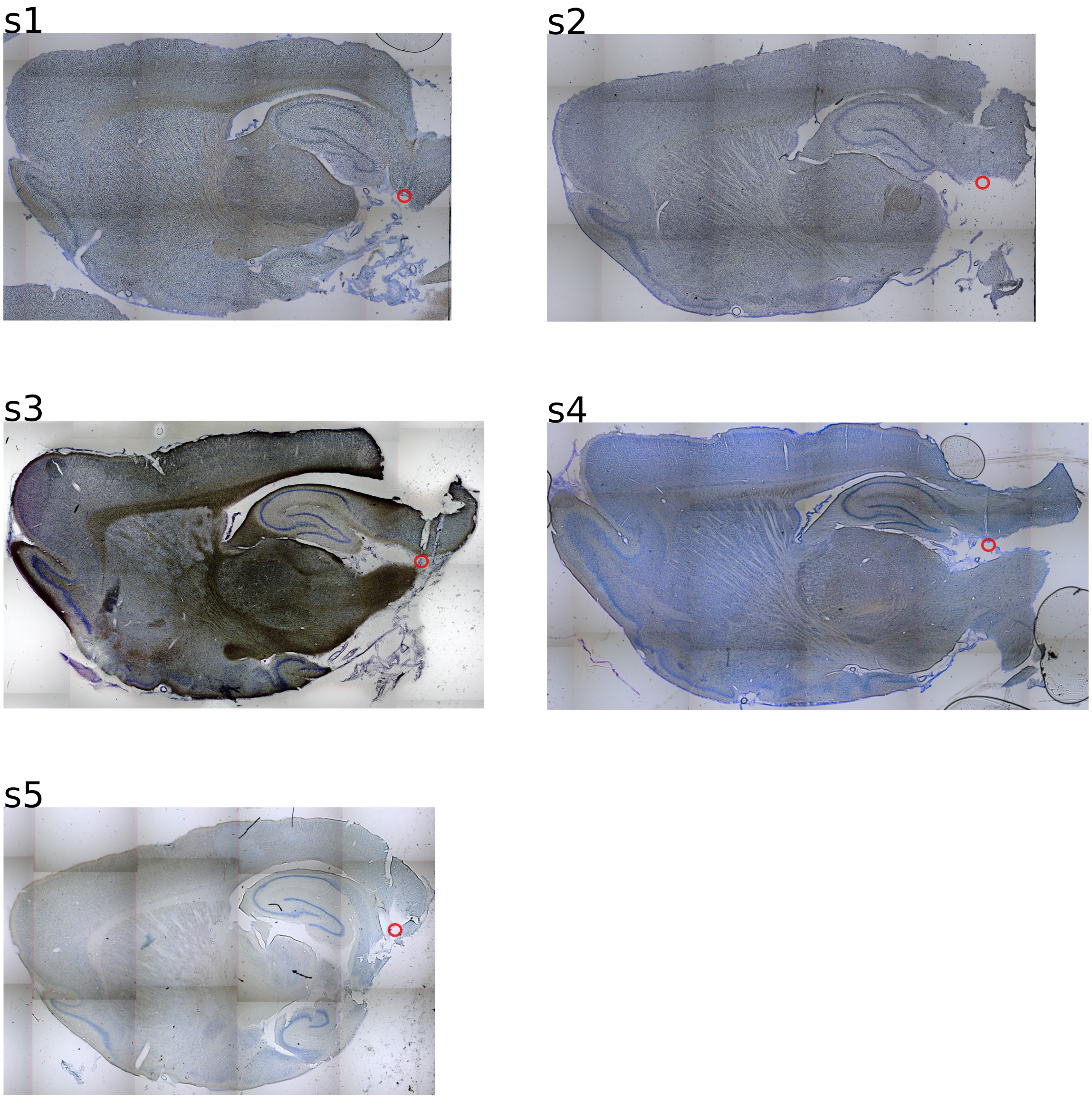


Supplemental Figure 3: Sagittal histology sections illustrating the location of recording electrodes. Coordinates for tetrode implantation ranged from 1 - 4 mm from the corital surface, 7.5 - 9 mm from bregma (anterior-posterior) and 3.6 - 4.6 mm from the midline (medial-lateral). Drive-able tetrodes with protective tubing were used and estimated end position highlighted with red circles. Tissue damage prohibited accurate localization of recording locations, with estimates being mostly MEC with some PaS and PrS.

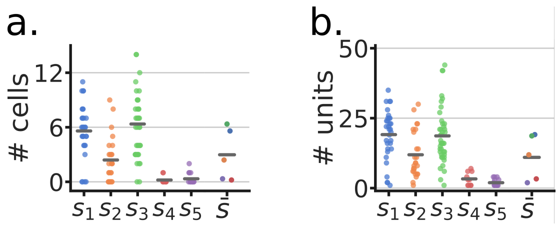


Supplemental Figure 4: Number of units by session and subject. **a.** Putative isolated units by subject. Each dot is a recorded session and horizontal bars indicate the mean by subject. The indication of $\bar{s}$ corresponds to subject means, color coded corresponding to each subject. **b.** Same as (**a**) but with the inclusion of MUA.

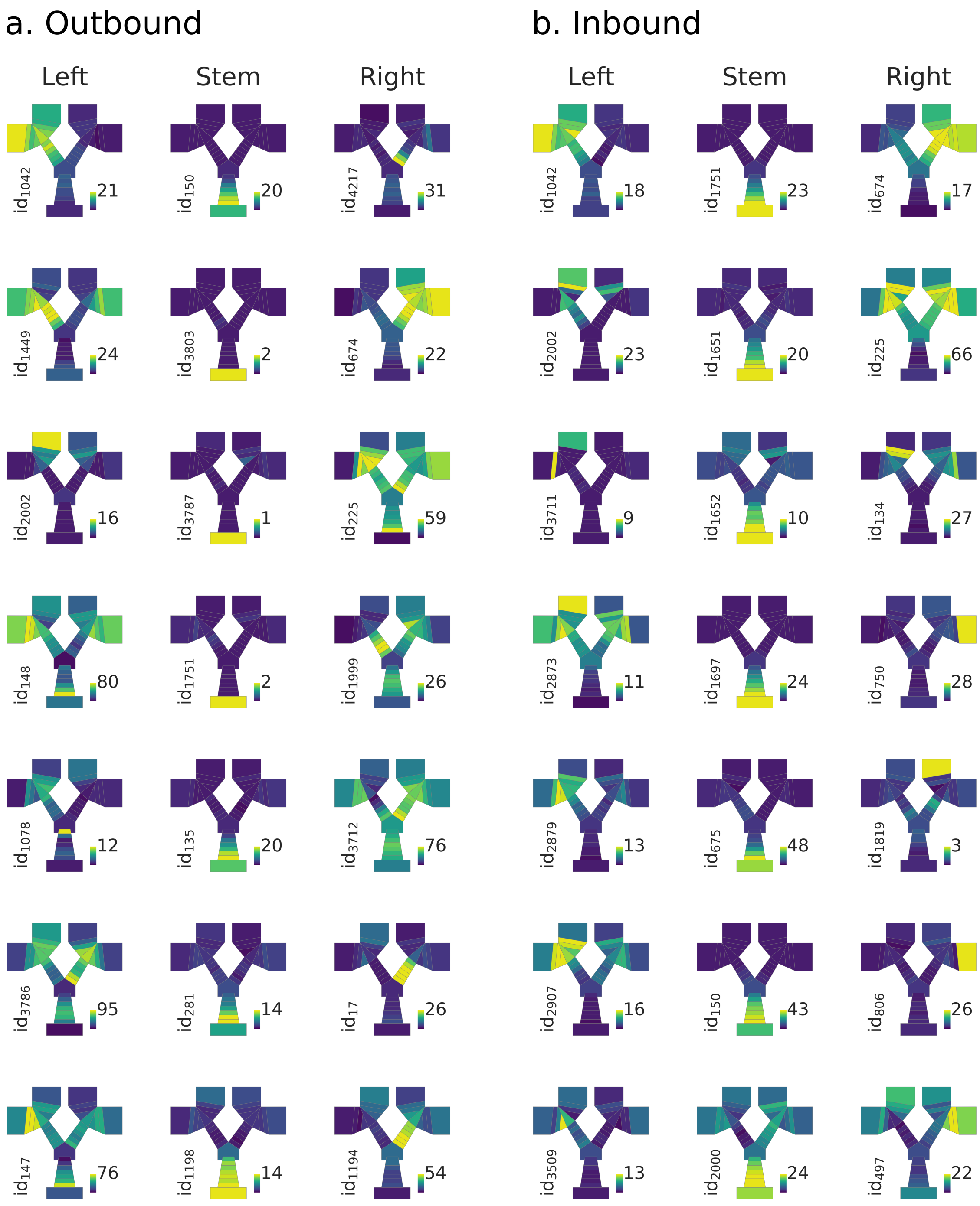


Supplemental Figure 5: Example rate maps of segment selective units. Units ranked (from top to bottom), by taking the Mann-Whitney $U_z$ statistic of that segment vs the two other segments (e.g. activity on Left vs activity on Stem and Right). **a-b.** Outbound trajectories (**a**) and inbound trajectories (**b**). Note that there is some overlap in selected units for inbound and outbound, indicating position tuning that was not directional.

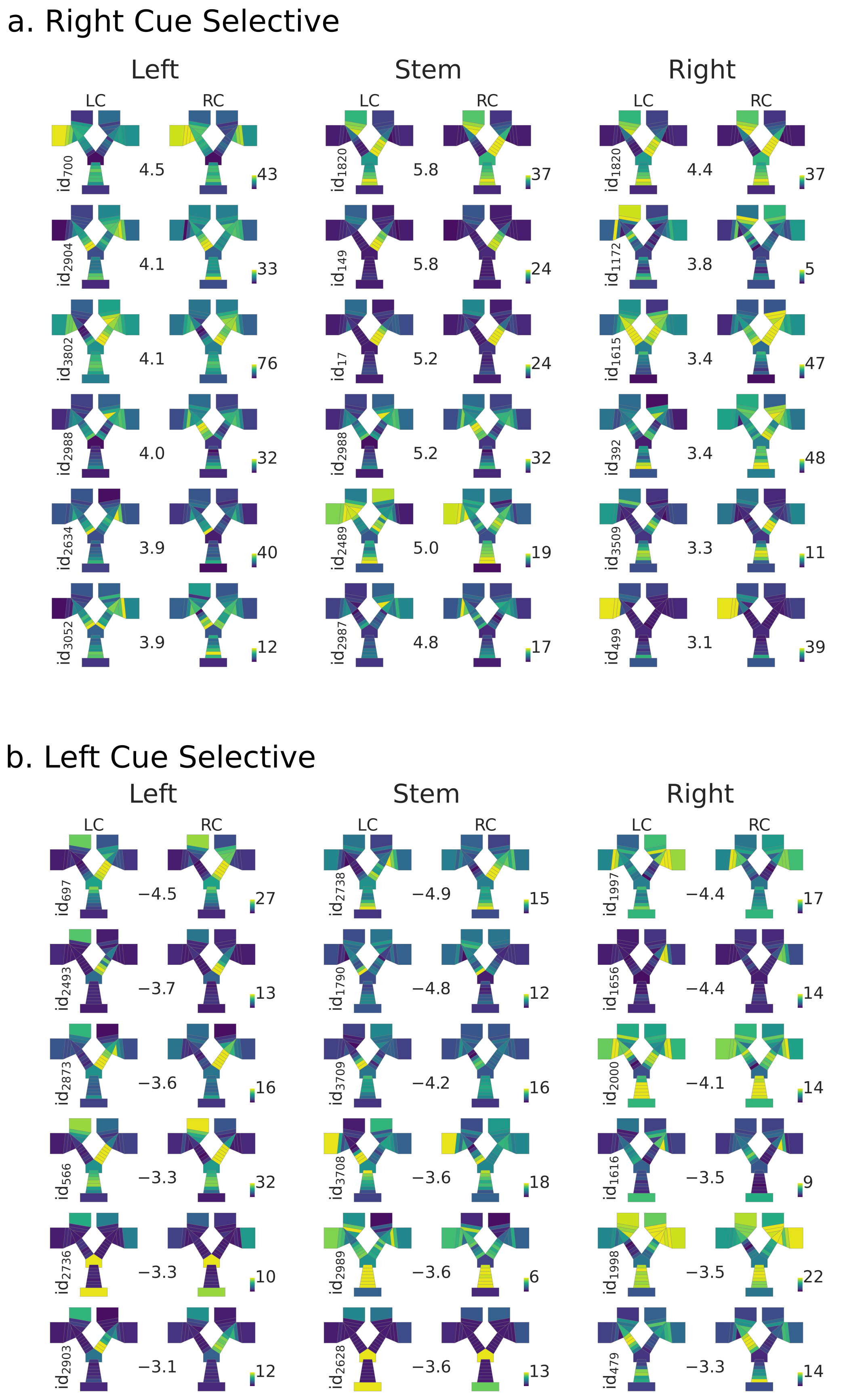


Supplemental Figure 6: Spatial rate maps of cue selective units. Units ranked (from top to bottom) by taking the Mann-Whitney $U_z$ statistic by segment of Right-Cue vs Left-Cue outbound trial conditions, statistic shown between each pair of rate maps. Columns refer to which segment of the maze was selected (left, stem, right). Rate maps are generated by averaging the activity by condition. Units shown were selected based on ranking of all the units according to the statistic value. The 90% of the max firing rate (spikes/second) of each pair of rate maps is shown in colormap, with all colormaps being referenced to 0 spikes/second. **a.** Spatial rate maps of right cue selective units by segment. Note that all $U_z$ values are positive. **b.** Spatial rate maps of left cue selective units by segment, all $U_z$ values are negative.

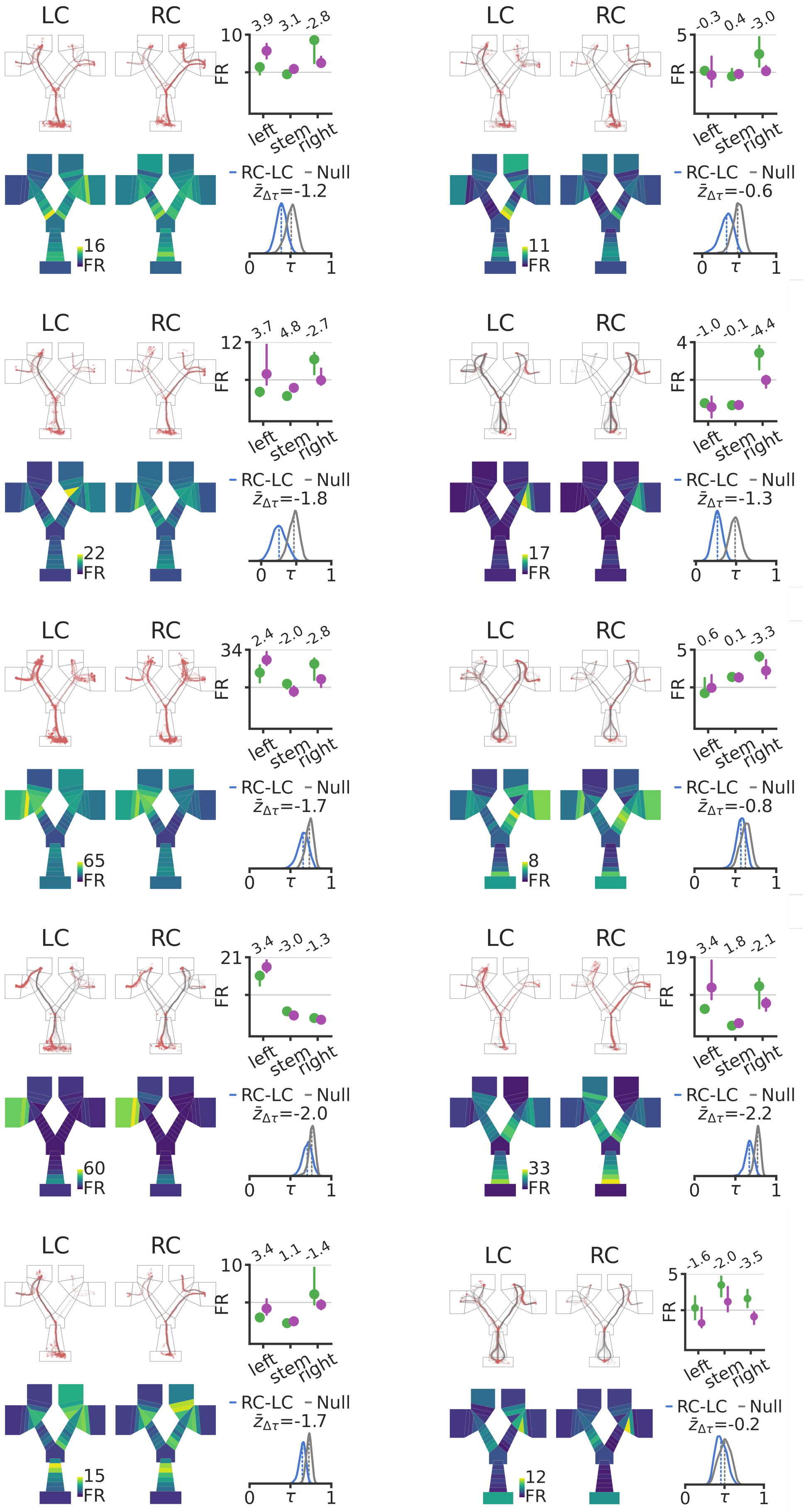


Supplemental Figure 7: Unit examples of stronger cue coding while in the incorrrect branch. Selection of units based on stronger coding for Left-Cue than Right-Cue when in the right branch of the maze, or stronger coding for Right-Cue than Left-Cue when in the left branch of the maze. Ten units, with each unit block showing raw spike/trajectories, rate maps, firing rate by segment x cue, and remapping distribution (as in Figure 2).

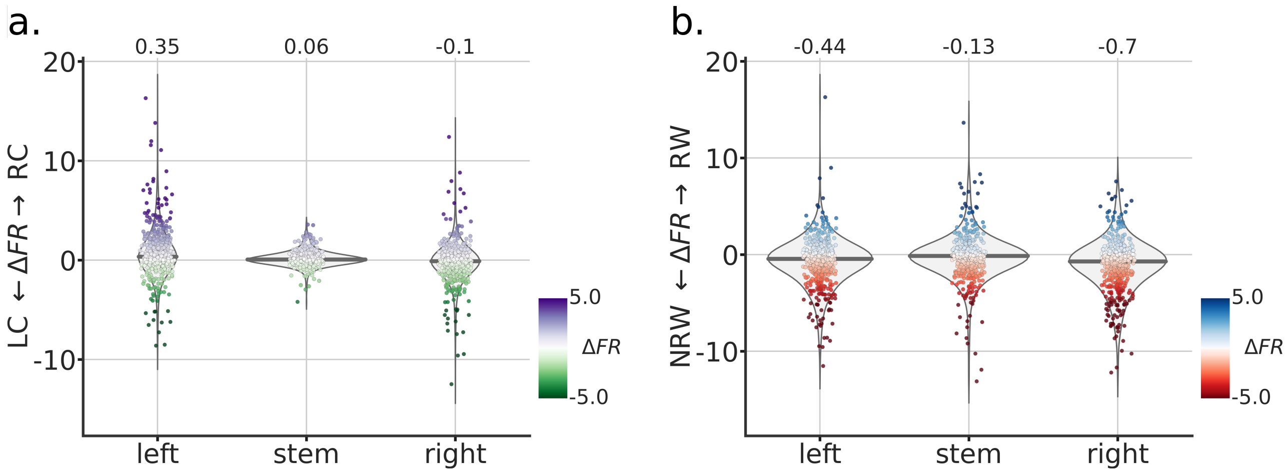


Supplemental Figure 8: Incorrect coding in mean firing rates. **a.** Plotted as in Figure 2c, with values being the difference of mean firing rates by cue condition and by maze segment. Even in this un-normalize space, the results follow what is reported in the main text. **b.** Plotted as in Figure 4c, with values being the difference of mean firing rates by reward condition and by maze segment.

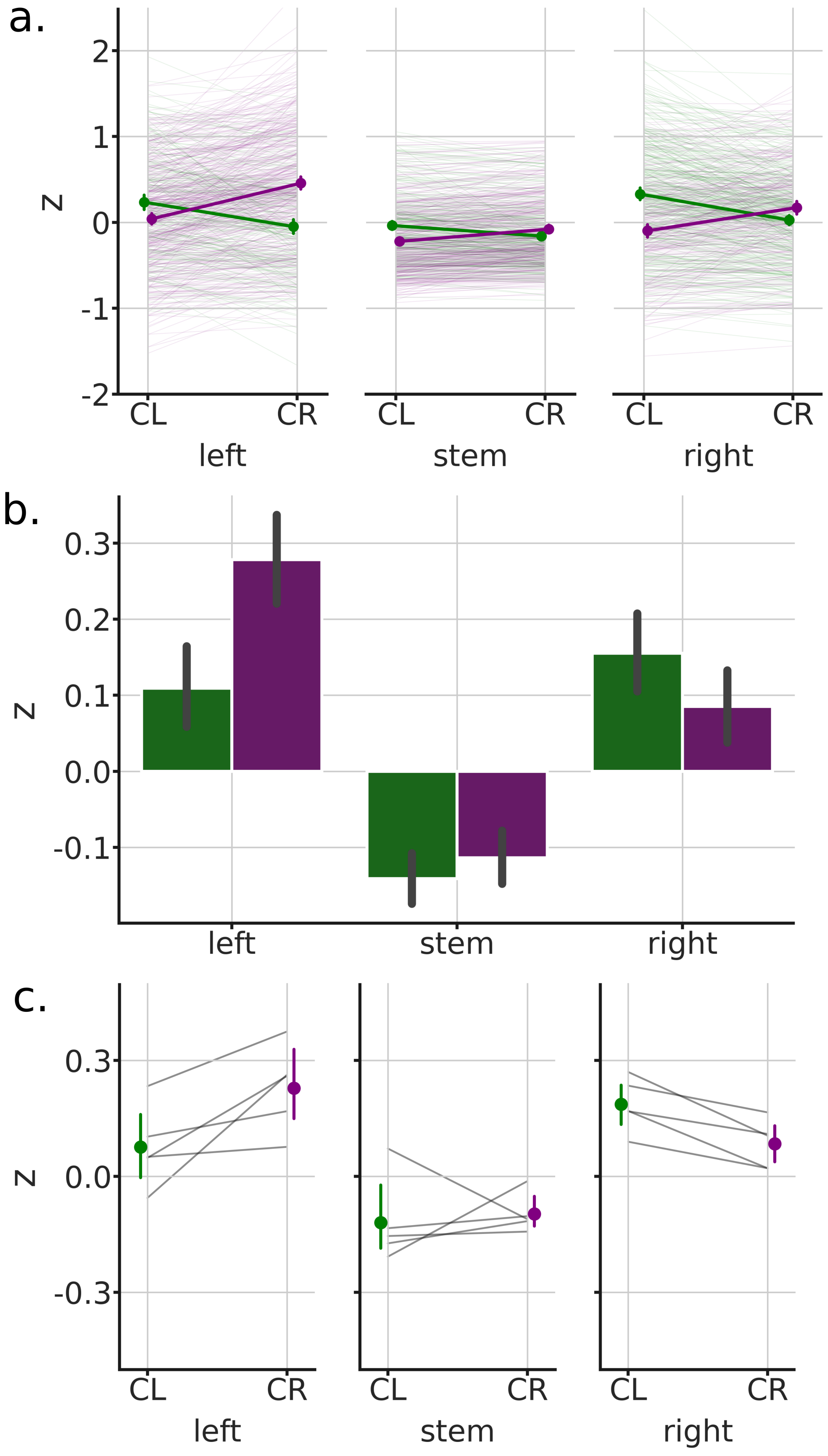


Supplemental Figure 9: Incorrect coding in Z-scored firing rates. **a.** Z-scored firing rate of all units by cue and maze segment. Individual unit color codes: Purple (CR>CL) and Green (CL>CR). Dot-lines are the averages for each of groupings. Note more "Purple" in the left segment of the maze and more "Green" in the right segment. Analysis is for illustration only as the pre-allocation of units to groups biases any analyses. **b.** Mean activity of all units by cue and maze segment. Note that all the general patterns of "error" / "incorrect" coding are as in the main analyses. **c.** Mean activity by subject. We replicate the above results by first taking the mean activity across each subject's units by cue and maze segment. Note that all the patterns are as in the main analyses.

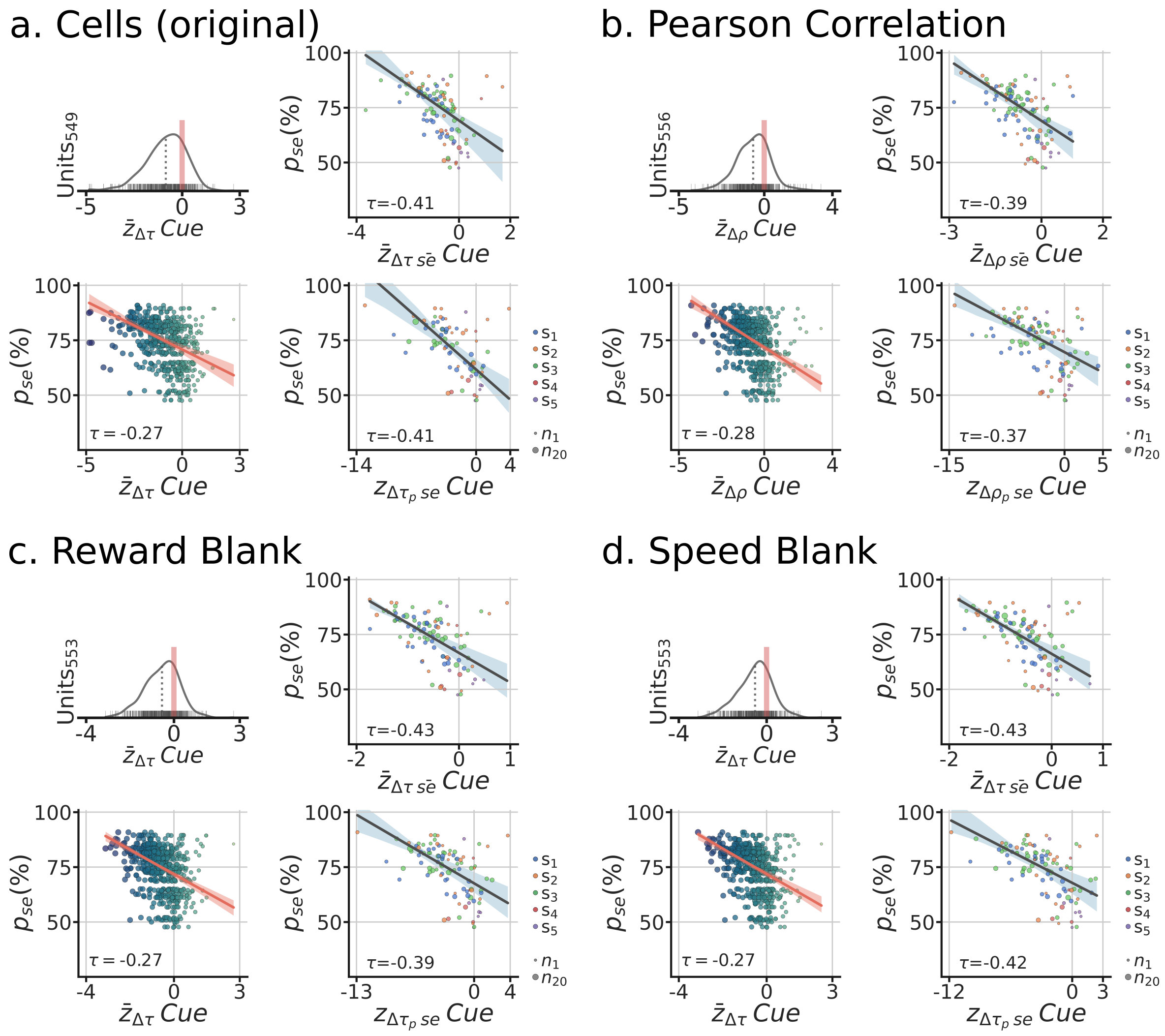


Supplemental Figure 10: Cue remapping vs behavior control analyses for isolated units. **a.** Original results as reported in Fig. 2. Top-left, distribution of remapping scores across units. Bottom-left, session performance $p_{se}$ by remapping score for each unit. Color and size of dots scale with x-axis. Red line is the robust regression line with 95% confidence band. Kendall $\tau$ between the quantities reported in the graph. Top-right, each dot is the average remapping score by session (color=subject, size= # of units in the average). Robust regression and 95% confidence band shown in dark blue. Bottom-right, each dot is the population correlation for the session. **b-d.** Positive control analyses, same as (a). **b.** Use of Pearson correlation instead of Kendall correlation to compute the remapping score. **c.** Each period post 500ms of reward delivery is removed from the analyses, avoiding possible artifacts due to activation of reward pump. **d.** Each period of speed being less than 2cm/s is removed from the analyses, only mobility periods go into the analyses. For all control analyses, we observed a significant relationship between remapping and behavior.

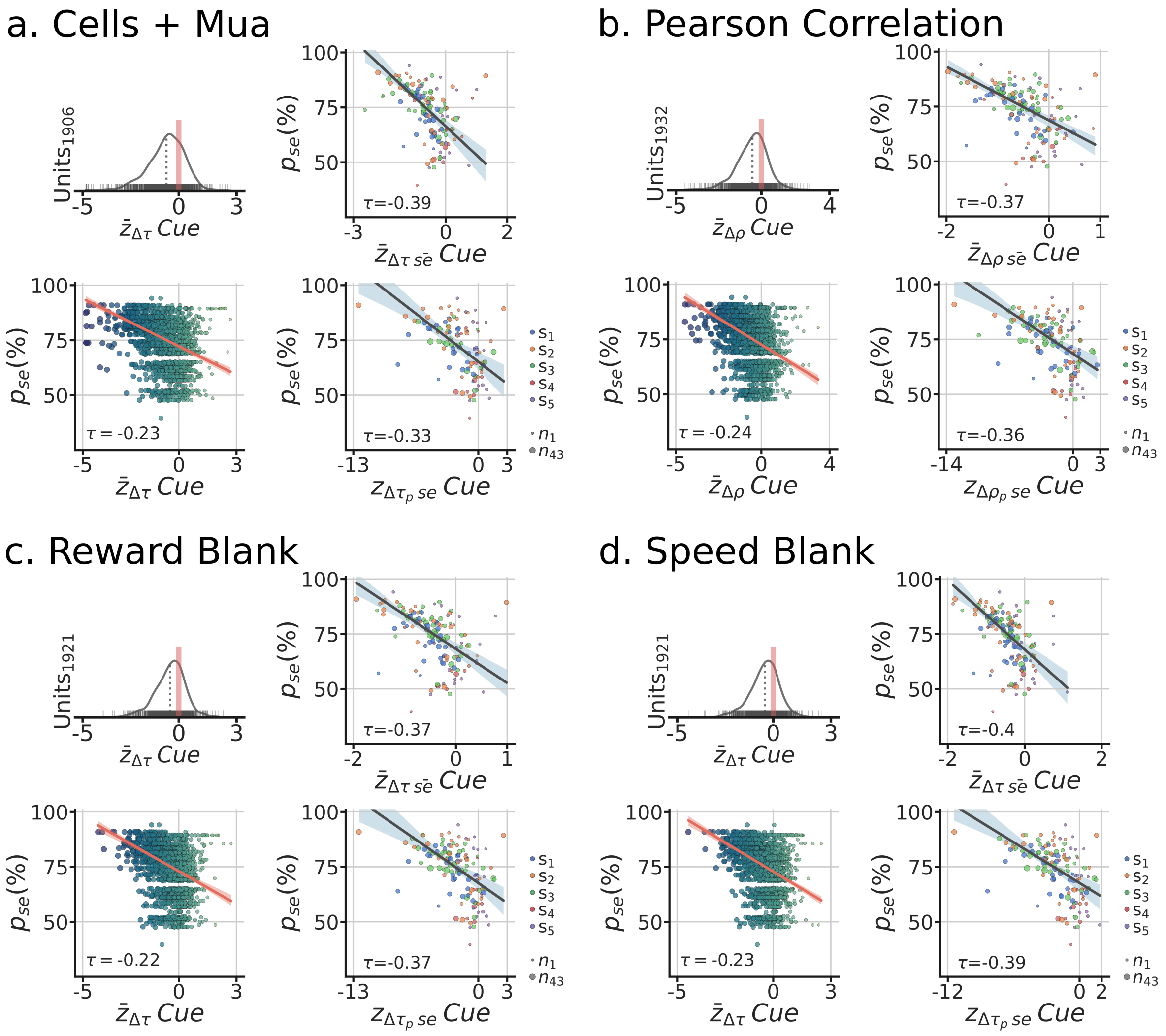


Supplemental Figure 11: Cue remapping vs behavior control analyses for isolated units and MUA. Panels follow convention in S.F.10. **a-d.** Top-left, distribution of remapping scores across units. Bottom-left, session performance $p_{se}$ by remapping score for each unit. Color and size of dots scale with x-axis. Red line is the robust regression line with 95% confidence band. Kendall $\tau$ between the quantities reported in the graph. Top-right, each dot is the average remapping score by session (color=subject, size= # of units in the average). Robust regression and 95% confidence band shown in dark blue. Bottom-right, each dot is the population correlation for the session (including both isolated units and MUA). **b.** Use of Pearson correlation instead of Kendall correlation to compute the remapping score. **c.** Each period post 500ms of reward delivery is removed from the analyses. **d.** Each period of speed being less than 2cm/s is removed from the analyses. Use For all control analyses, we observed a significant relationship between remapping and behavior.

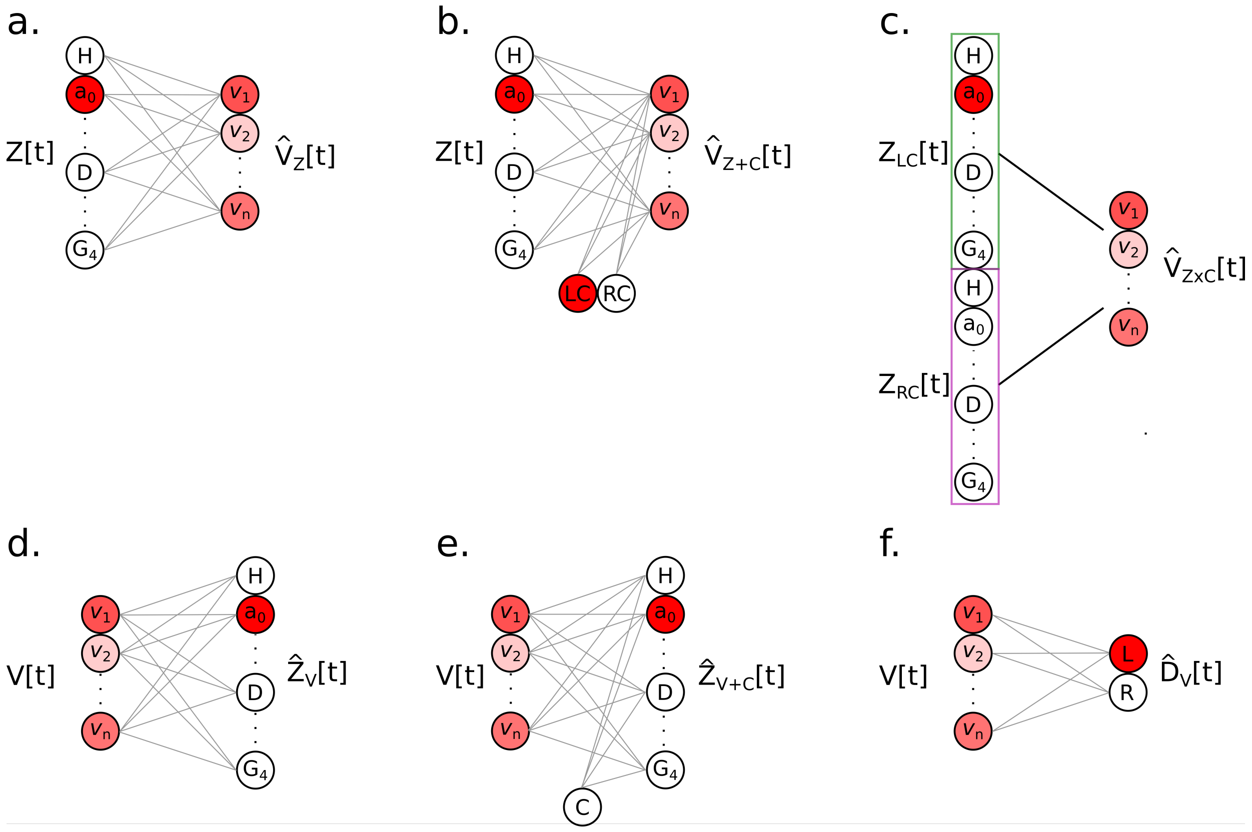


Supplemental Figure 12: Encoder and decoder model diagrams. **a.** Zone encoder model, $Z$, that uses the position of the animal at time [t] to predict the neural activity $V$ after a linear transformation. Note that only one zone can be active at a time. Intensity of red color indicates activity. **b.** Zone encoder plus cue identity model ($Z + C$, rate-remapping model). Note that only one cue can be active by trial, effectively functioning as a single modulatory gain signal that would change the activity across the trial duration. **c.** Zone encoder by cue identity model ($ZxC$, global-remapping model). This model had 2 sets of positions, with only one set active as a function of cue, effectively allowing for completely orthogonal maps to be used to predict the neural activity. Position sets indicated by color. Reward encoding models followed the same convention, with RW instead of cue. **d.** Zone decoder model that uses the neural activity to predict the subject's position, $V_Z$. Omitted from the diagram is the presence of a softmax normalization that produces probability values for each zone. The maximum probability at a given time is used as the prediction of position for that time. **e.** Zone decoder model, like (**d**), plus the inclusion of cue information $V_{Z+C}$. Note that a weighted cue value is added to neural activity by zone, before the softmax operation. This approach thus allow the cue to have different effects by zone. **f.** Decision decoder, $D$, that uses neural activity to predict the subject's decision (Left/Right) by time. Omitted from the diagram is the cumulative temporal integration of predictions by trial, allowing the decoder to accumulate evidence as the subject traverses the maze.

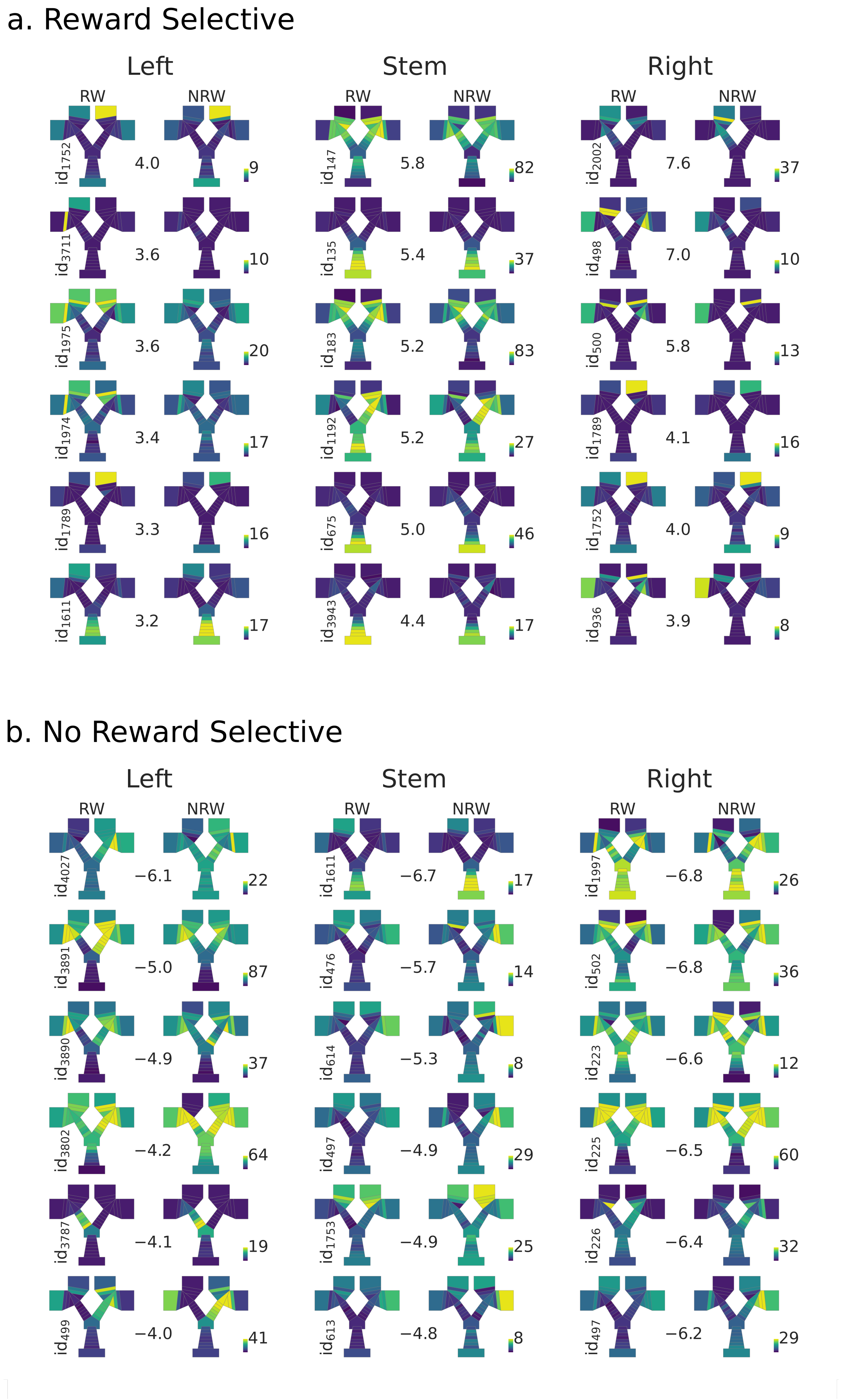


Supplemental Figure 13: Spatial rate maps of reward selective units. Units ranked (from top to bottom) by taking the Mann-Whitney $U_z$ statistic by segment of reward vs no-reward inbound trials conditions, statistic shown between each pair of rate maps. Columns refer to which segment of the maze was selected (left, stem, right). Rate maps are generated by averaging the activity by condition. Units shown were selected based on ranking of all the units according to the statistic value. The 90% of the max firing rate (spikes/second) of each pair of rate maps is shown in colormap, with all colormaps being referenced to 0 spikes/second. **a.** Spatial rate maps of reward selective units by segment. Note that all $U_z$ values are positive. **b.** Spatial rate maps of no-reward selective units by segment, all $U_z$ values are negative.

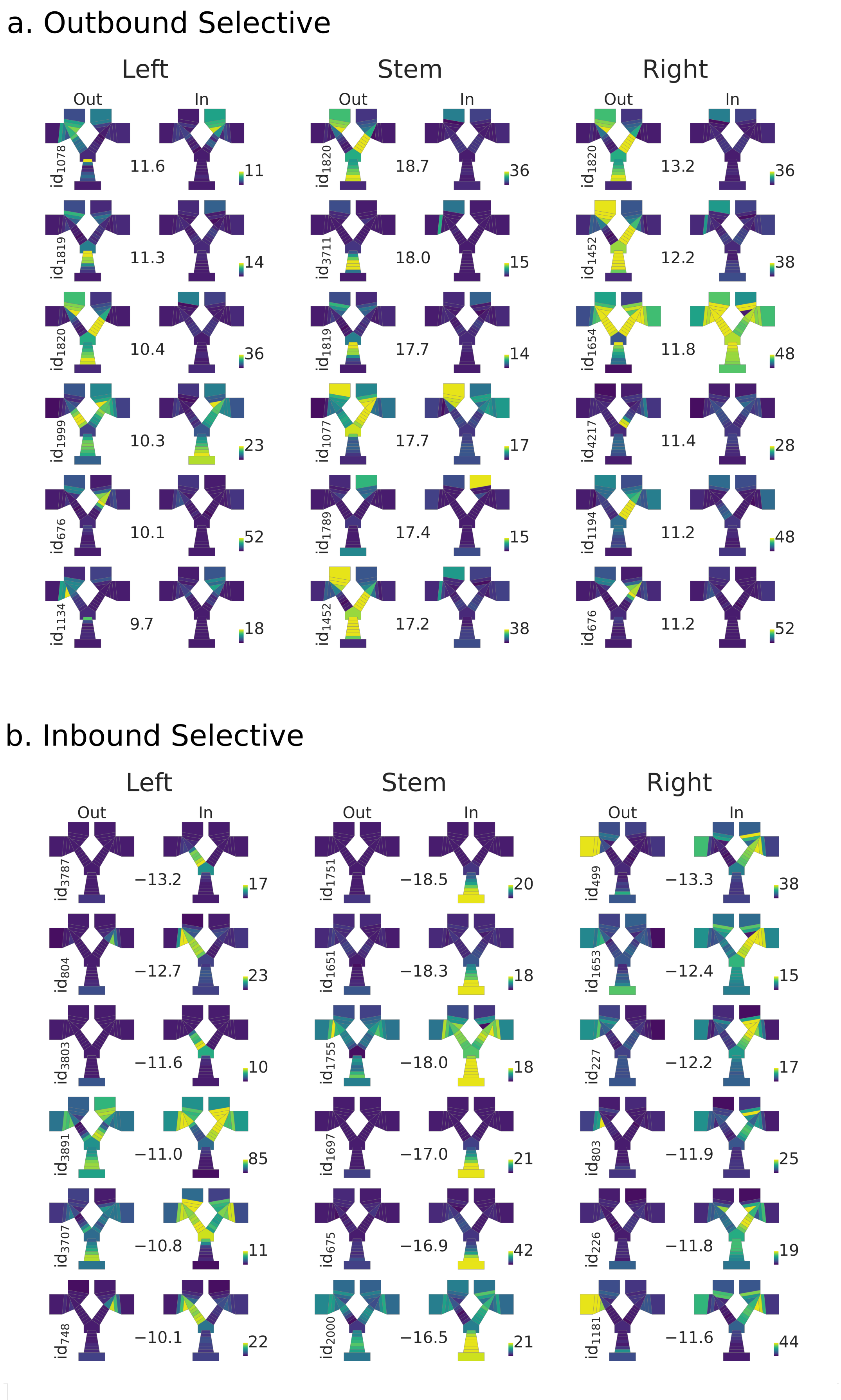


Supplemental Figure 14: Spatial rate maps of direction selective units. Units ranked (from top to bottom) by taking the Mann-Whitney $U_z$ statistic by segment of outbound vs inbound trajectories for all trials, statistic shown between each pair of rate maps. Rate maps are generated by averaging the activity by condition. Units shown were selected based on ranking of all the units according to the statistic value. The 90% of the max firing rate (spikes/second) of each pair of rate maps is shown in colormap, with all colormaps being referenced to 0 spikes/second. **a.** Spatial rate maps of outbound selective units by segment. Note that all $U_z$ values are positive. **b.** Spatial rate maps of inbound selective units by segment, all $U_z$ values are negative.

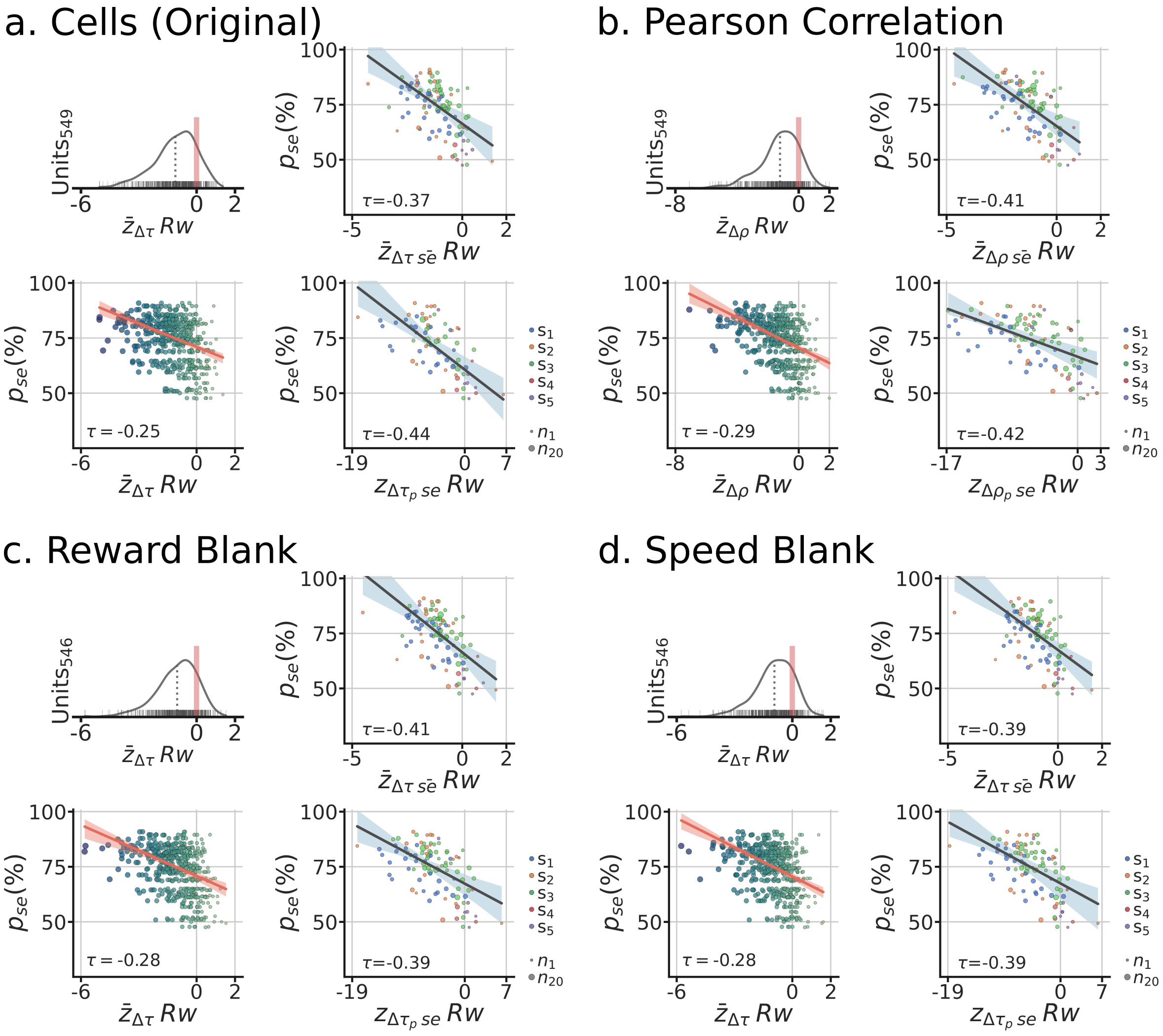


Supplemental Figure 15: Reward remapping vs behavior control analyses for isolated units. Panels follow convention in S.F.10. **a-d.** Top-left, distribution of remapping scores across units. Bottom-left, session performance $p_{se}$ by remapping score for each unit. Color and size of dots scale with x-axis. Red line is the robust regression line with 95% confidence band. Kendall $\tau$ between the quantities reported in the graph. Top-right, each dot is the average remapping score by session (color=subject, size= # of units in the average). Robust regression and 95% confidence band shown in dark blue. Bottom-right, each dot is the population correlation for the session. **b.** Use of Pearson correlation instead of Kendall correlation to compute the remapping score. **c.** Each period post 500ms of reward delivery is removed from the analyses. **d.** Each period of speed being less than 2cm/s is removed from the analyses. For all control analyses, we observed a significant relationship between remapping and behavior.

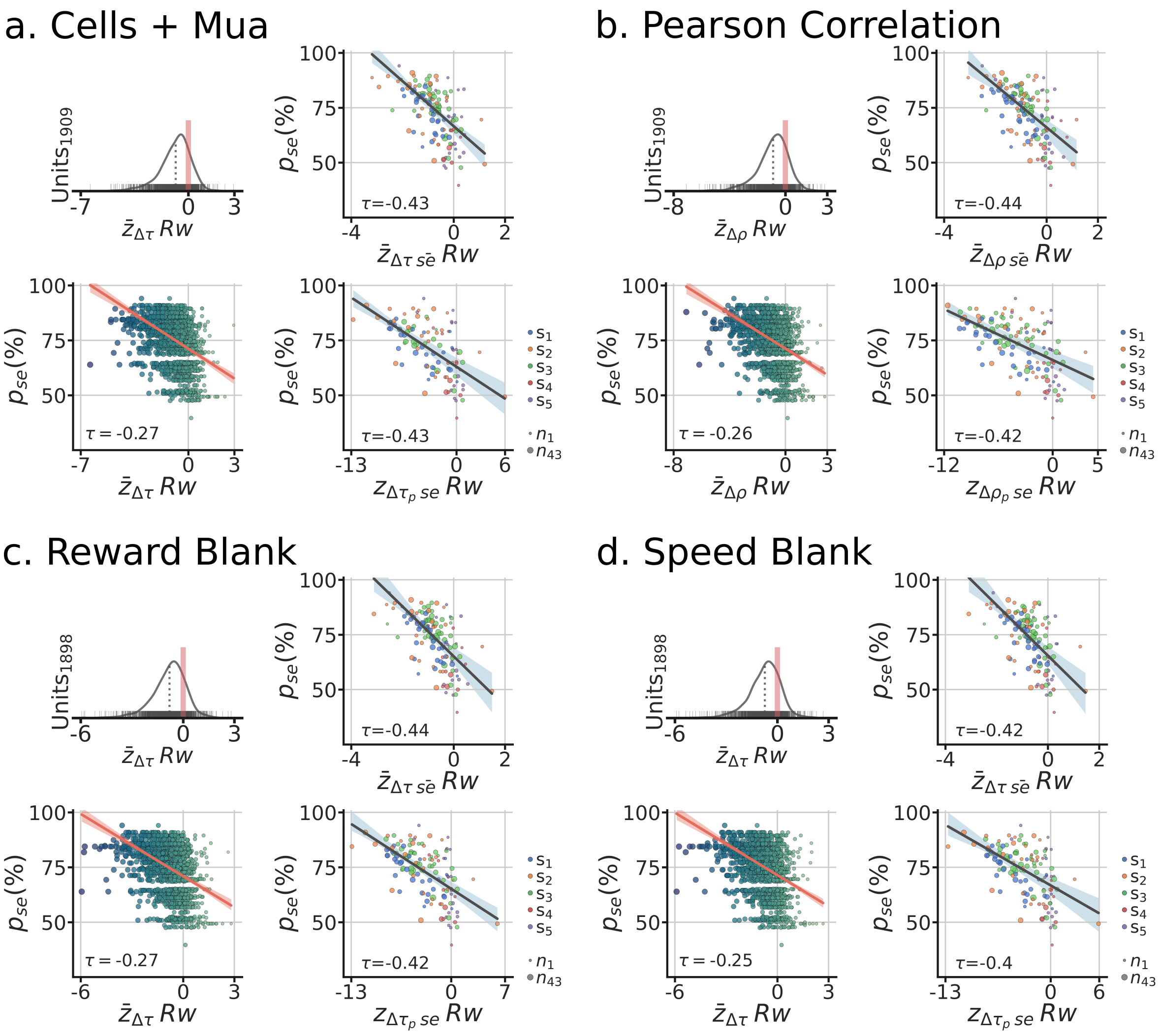


Supplemental Figure 16: Reward remapping vs behavior control analyses for isolated units and MUA. Panels follow convention in S.F.10. **a-d.** Top-left, distribution of remapping scores across units. Bottom-left, session performance $p_{se}$ by remapping score for each unit. Color and size of dots scale with x-axis. Red line is the robust regression line with 95% confidence band. Kendall $\tau$ between the quantities reported in the graph. Top-right, each dot is the average remapping score by session (color=subject, size= # of units in the average). Robust regression and 95% confidence band shown in dark blue. Bottom-right, each dot is the population correlation for the session (including both isolated units and MUA). **b.** Use of Pearson correlation instead of Kendall correlation to compute the remapping score. **c.** Each period post 500ms of reward delivery is removed from the analyses. **d.** Each period of speed being less than 2cm/s is removed from the analyses. For all control analyses, we observed a significant relationship between remapping and behavior.

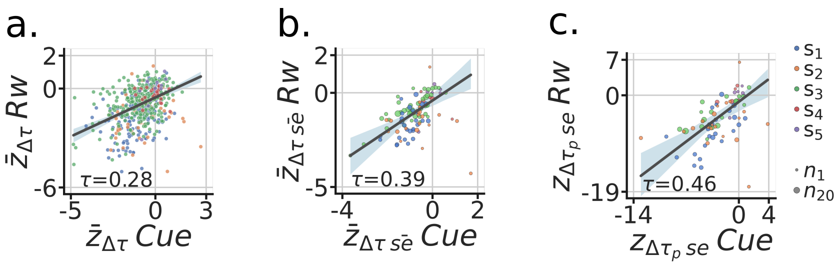


Supplemental Figure 17: Relationship between remap scores for cue and reward. **a.** Remap scores for each unit, colored by subject identity. **b.** Mean remap scores by session, colored by subject identity and size indicates number of units. **c.** Population correlation by session. Note that remapping scores correlate across the three levels of analyses, suggesting circuit coherence throughout the behavioral distinct outbound/inbound segments of a trial.

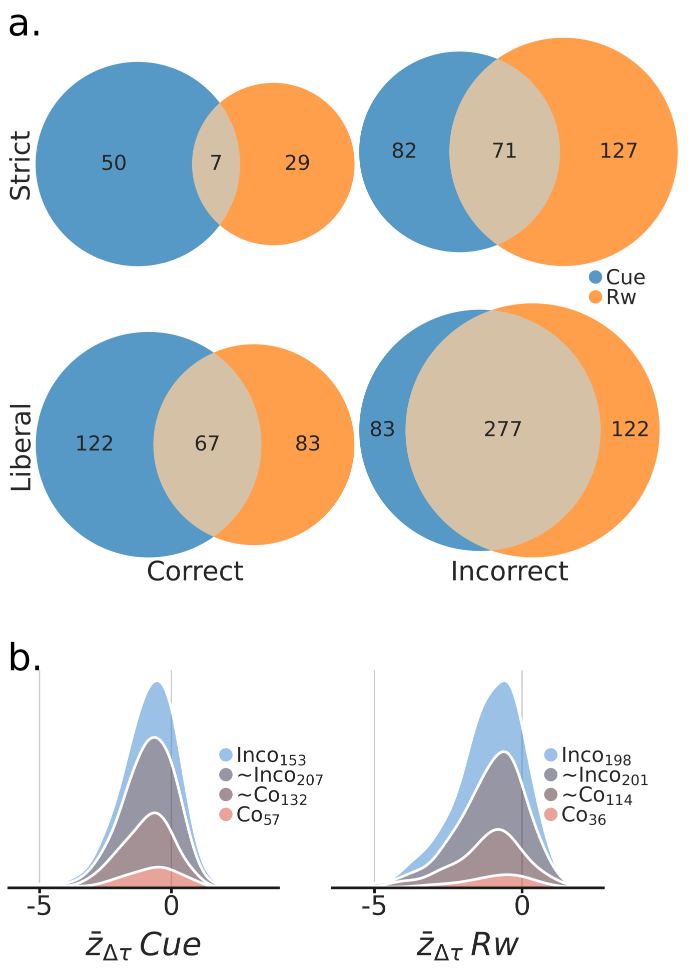


Supplemental Figure 18: Cue and reward rate coding vs correct/incorrect interaction. For these analyses, the patterns of cue coding $U_{Z_{RC-LC}}$ and reward coding $U_{Z_{RW-NRW}}$ by segment and unit were used to determine correct/incorrect coding (Methods). **a.** Venn diagrams for the overlap between cue/reward coding units by correct/incorrect coding. Top-row, strict correct/incorrect coding criteria, Bottom-row, liberal correct/incorrect coding criteria (super-set of top-row). Strict criteria required correct (incorrect) coding present in all maze segments (3 out of 3 segments), while liberal criteria classification was based on 'most' segments (2 out 3 Segments). Columns correspond to correct and incorrect, and colors to cue or reward. Note that the overlap in coding largely comes from incorrect coding. Overlap of incorrect coding between cue and reward is greater than the overlap in correct (Fisher's Exact Test $p = 0.003$ for strict, $p = 0.007$ for liberal). **b.** Distributions of unit remapping scores by the different groupings of correct/incorrect coding and cue/reward. Qualitatively, correct/incorrect coding was not predictive of the units remapping strength (all Kolmogorov-Smirnov 2 sample tests $p > 0.17$). [Co=correct coding, Co=weakly codes for correct, Inco=weakly codes for incorrect, Inco=incorrect coding].

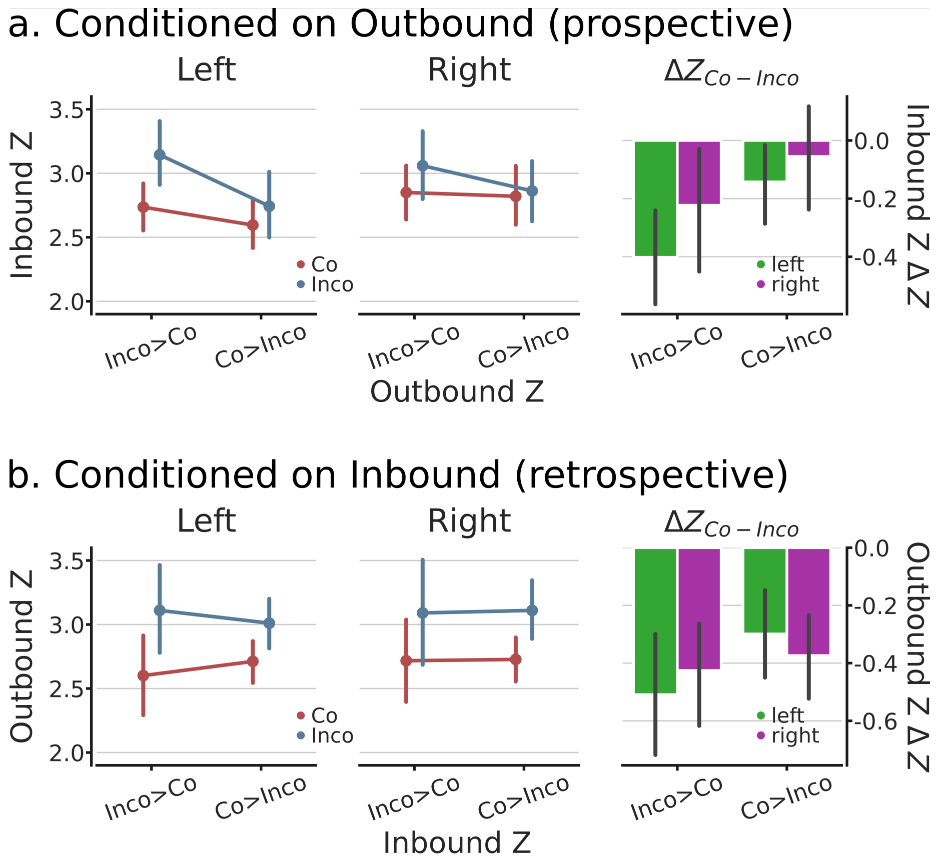


Supplemental Figure 19: Correct vs incorrect rates during the transition between Outbound and Inbound (return) trajectories. Analyses on this figure are performed on z-scored mean firing rates for correct (Co) and incorrect (Inco) trials (minimum of 5 Inco trials in the session) for single units. **a.** Conditioned on Outbound rate differences between Co and Inco, the Inbound activity are overall larger for incorrect trials (Left and Middle panels, reflecting the Left and Right branches of the maze; LMEM: main effect of Inco $LRT = \chi^2_1 = 6.64$, $p = 9.97e^{-3}$). Right panel, by unit differences between Co and Inco, showing that if Inco>Co on the Outbound, then Inco>Co on the Inbound trajectory. The Co>Inco did not show this effect (LMEM: interaction $LRT = \chi^2_1 = 6.89$, $p = 8.67e^{-3}$). This observations implies that, on average and across units, elevated activity levels during incorrect trials on the Outbound trajectories are sustained during the subsequent Inbound trajectories, while higher activity rates on Outbound correct trajectories do not show any predictability on the subsequent inbound activity. **b.** Conditioned on Inbound rates differences between Co and Inco, preceding Outbound rates were overall higher for incorrect trials (LMEM: main effect of Inco $\chi^2_1 = 24.28$, $p = 8.34e^{-7}$). Right panel are the differences between Co and Inco on the Outbound rates, showing that the reverse inference is also significant (LMEM: interaction $\chi^2_1 = 9.67$, $p = 1.93e^{-3}$). All LMEM had fixed effects of segment (Left, Right), the conditioning variable (Inco>Co, Co>Inco), and for the main effect LMEMs condition (Inco, Co). Additionally, subjects were modeled as a random effect, with additional variance components of task version and session. Summarizing, Inco>Co on Outbound trajectories predicts that the subsequent Inbound trajectories will also have Inco>Co, similarly Inco>Co on Inbound trajectories predict that the preceding Outbound trajectory rates also were Inco>Co.

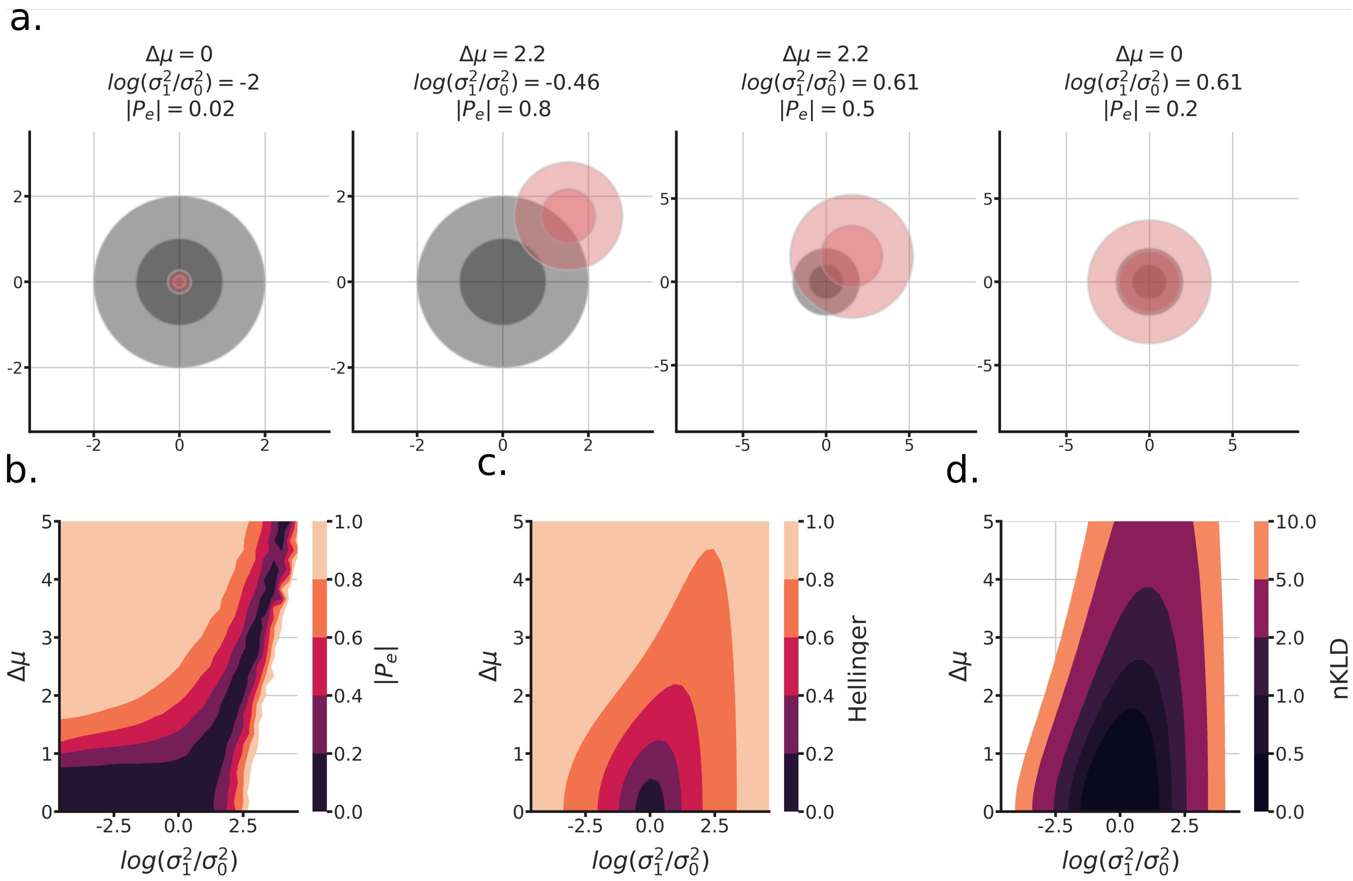


Supplemental Figure 20: Demonstration of waveform matching algorithm. This algorithm was followed to determine which units were recorded in both the open-field and Tree-Maze tasks. We operationalized this matching by quantifying how likely it is that given the waveform of random spike it would be incorrectly assigned to the correct cluster. **a.** Four simulated scenarios of the resulting wave clusters for two units, parameterized by their difference in location $\Delta\mu$, and their log-ratio of variances $log(\frac{\sigma_1^2}{\sigma_0^2})$. The metric $|P_e|$ is also shown for each example, this represents the probability of mistaking a waveform for the other. If this value is less than 0.5, we say that given the data, those units are not distinguishable from another and are thus matched. In these examples, only the second from the left would not be matched $|P_e| = 0.8$. Note that while asymmetry is possible, (e.g. $P_e$ of $unit_0$ to $unit_1$ is different than $P_e$ of $unit_1$ to $unit_0$), that scenario was not of interest for our study and $P_e$ values are averaged for each pair. **b.** Full landscape of $P_e$ for the parametrization between a pair of units. Note that there is a band of matching along this landscape. **c-d** Hellinger and normalized KLD distances. Note that these metrics tend to reflect the "closeness" of the mean/variance of the distributions, which is not the main goal of the matching algorithm.

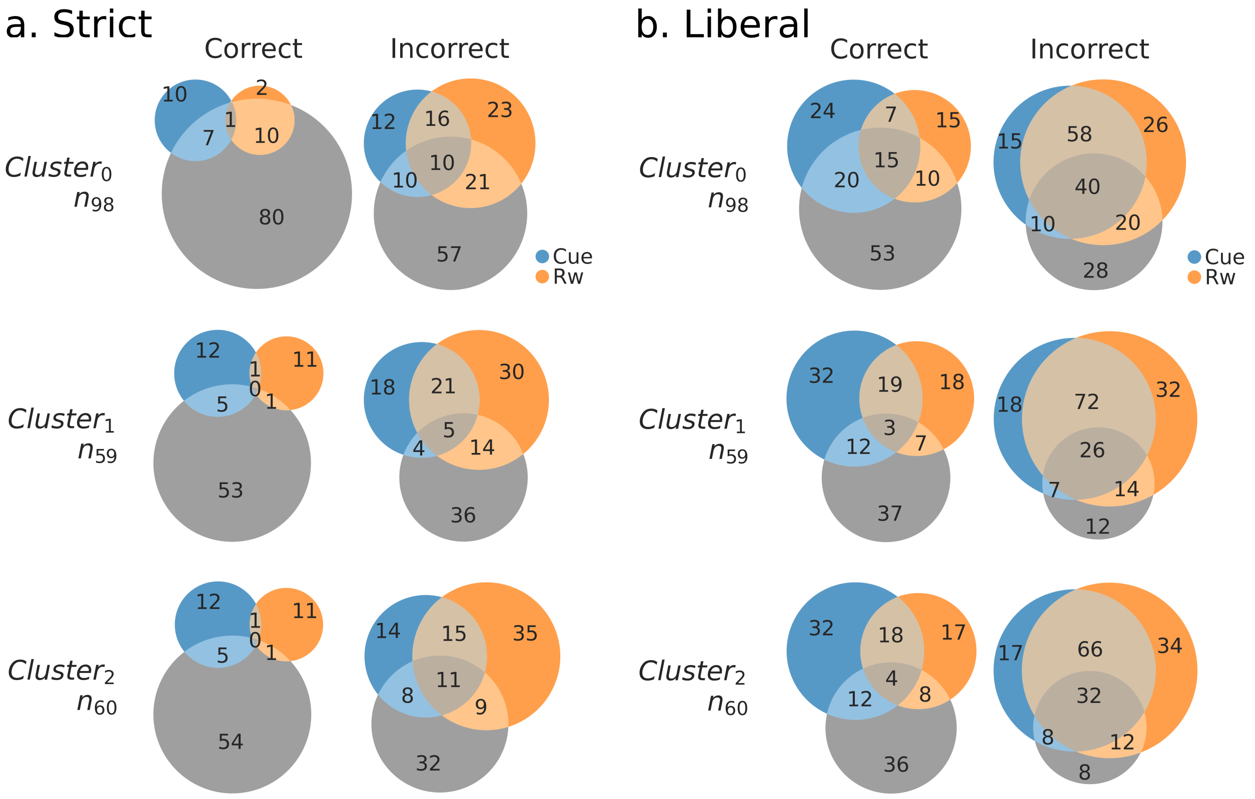


Supplemental Figure 21: Cue and reward rate coding vs correct/incorrect interaction and cluster identity. Functionally defined clusters of units identified in the open-field. Thresholds for correct/incorrect as described in 18. **a.** Strict correct/incorrect coding. **b.** Liberal correct/incorrect coding, groupings in **a.** are part of **b.**. There is little overlap between the open-field clusters and correct coding for both cue and reward, while the overlap is larger for incorrect coding. At the same time, most incorrect coding units did not fall into any of the clusters, suggesting that the Tree-Maze task recruits a different population of neurons that are either inactive or were not detected in the Open-Field task.

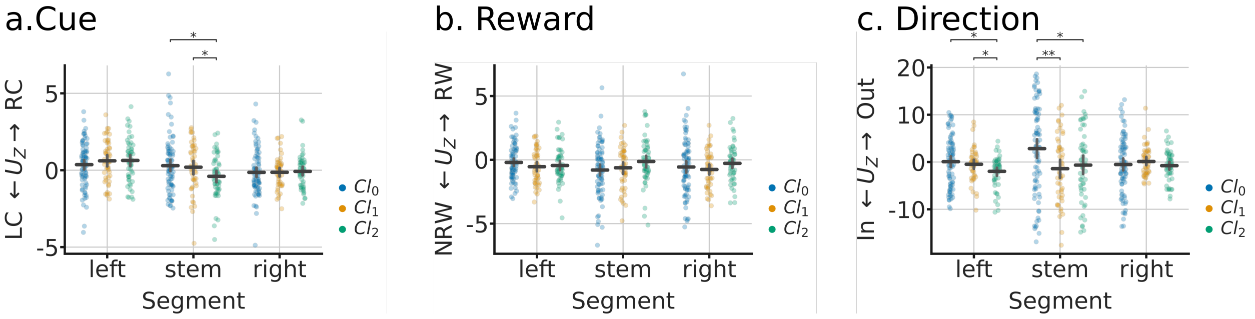


Supplemental Figure 22: Differences in activity rates by cluster and condition. These analyses use the functionally defined clusters of units identified in the open-field arena. **a.** Outbound cue firing rate activity difference (Mann-Whitney statistic $U_Z$) between Right-Cue and Left cue by segment and cluster. **b.** Inbound differences between rewarded (RW) and not-rewarded (NRW) trials. **c.** Differences in representation for Outbound (Out) and Inbound (In) trajectories. Note the higher variability for cluster 0, consistent with that cluster containing Head-Direction units.
